# TDP-43 subtypes shape transcriptomic signatures in Alzheimer’s disease

**DOI:** 10.64898/2026.06.22.733827

**Authors:** Xiaojie Wang, Alyssa C. Walker, Madison M. Reeves, Jesus Gonzalez Bejarano, Yuping Song, Judy Dunmore, Mei Yue, Bailey Rawlinson, Erica Engelberg-Cook, Michael DeTure, Raul Baena Tome, Anand Narayan, Jordan Bartfield, Neill R. Graff-Radford, Bradley F. Boeve, Ronald C. Peterson, David S. Knopman, Björn Oskarsson, Gregory S. Day, Melissa E. Murray, Dennis W. Dickson, Casey Cook, Yongjie Zhang, Leonard Petrucelli, Keith A. Josephs, Mercedes Prudencio

## Abstract

TAR DNA-binding protein 43 (TDP-43) pathology frequently co-occurs with Tau neurofibrillary tangles (NFTs) and amyloid β plaques in Alzheimer’s disease (AD), driving significant clinical heterogeneity. Whether TDP-43 engages autonomous molecular programs or instead amplifies Tau-driven neurodegeneration remains difficult to resolve, largely because these pathologies often co-occur. To separate these overlapping signatures, we generated regionally resolved transcriptomic profiles from cognitively normal controls (Controls), neuropathologically defined cohorts of AD, AD with limbic-predominant age-related TDP-43 encephalopathy (AD/LATE), and frontotemporal lobar degeneration (FTLD-TDP), categorizing them by their distinct TDP-43 subtypes (types α and β for AD/LATE; types A and B for FTLD-TDP). By integrating transcriptomic profiles with quantitative measures of phosphorylated TDP-43 (pTDP-43) and Tau (pTau), we separated pathology-associated signals within mixed disease contexts. We found that TDP-43 is linked to distinct transcriptomic programs in AD/LATE that are largely uncoupled from Tau burden and diverge from those observed in FTLD-TDP. These signatures showed regional specificity, with transcriptomic remodeling occurring in the amygdala across both diseases, whereas frontal cortex alterations were largely restricted to FTLD-TDP. Furthermore, by stratifying cases by TDP-43 morphological subtype, we unmasked specific biological trajectories, from immune activation to unique cellular vulnerabilities, that are not apparent in unstratified cohorts. Together, our findings provide a framework for decoupling mixed proteinopathies and demonstrate that TDP-43 shapes autonomous, subtype-dependent transcriptional landscapes in AD.

## Main

Alzheimer’s disease (AD) is neuropathologically defined by the accumulation of amyloid-β plaques and Tau neurofibrillary tangles (NFTs). However, these pathologies do not fully account for the extreme variance in clinical outcomes^1–3^. Much of this heterogeneity stems from comorbid proteinopathies, particularly TAR DNA-binding protein 43 (TDP-43)^2–4^. Beyond its role in frontotemporal lobar degeneration (FTLD-TDP) and amyotrophic lateral sclerosis (ALS) ^5,6^, TDP-43 is a frequent co-pathology in AD and the aging brain, where it is termed limbic-predominant age-related TDP-43 encephalopathy (LATE)^2–4^. The presence of TDP-43 pathology in AD (AD/LATE) significantly accelerates cognitive decline and accounts for a more severe clinical trajectory^1–4,7^, suggesting that TDP-43 contributes independently to disease progression.

Despite increasing recognition of TDP-43 as an important co-pathology in AD, defining its specific molecular effects has remained challenging because it almost invariably occurs in the context of established AD pathology. TDP-43 pathology in AD/LATE is largely confined to limbic structures, frequently coexisting with tau, while amyloid-β and tau pathologies spread extensively throughout the neocortex^3,4,8^.

Similar to the classical TDP-43 subtypes in FTLD-TDP (types A–E)^9,10^, TDP-43 pathology in AD/LATE is commonly classified into two distinct subtypes: type α, characterized by cytoplasmic inclusions and abundant dystrophic neurites; and type β, defined by diffuse or fibrillary inclusions that co-mingle with NFT^11,12^.

To identify disease-specific and TDP-43-specific changes in AD, we performed bulk RNA sequencing in amygdala and frontal cortex tissues from a cohort consisting of cognitively normal controls, AD without TDP-43 pathology (AD no TDP), AD/LATE, and FTLD-TDP. AD/LATE and FTLD-TDP cases were further stratified by morphological subtype (type α and β for AD/LATE; and the most abundant type A and B for FTLD-TDP). We also integrated transcriptomic profiles with quantitative biochemical measures of insoluble phosphorylated TDP-43 (pTDP-43) and Tau (pTau) to identify gene sets and molecular pathways associated with each pathology. Together, our findings define molecular features of TDP-43 proteinopathies in the context of AD and provide a framework for distinguishing TDP-43-associated signatures from Tau-driven neurodegeneration.

## Results

### Neuropathological features of AD and FTLD-TDP cohorts

To investigate how TDP-43 dysfunction contributes to transcriptomic changes specific to AD, we performed RNA sequencing on postmortem brain tissues from neuropathologically-confirmed AD/LATE (N=59) and AD no TDP (N=21), controls (N=22), and FTLD-TDP cases (N=56) (**Fig.1a**, **Table 1, Extended Data Table 1**). All AD cases exhibited advanced AD pathology, with median Braak stage of 6 and a median Thal stage of 5 (**Table 1**). The prevalence of comorbid vascular dementia was comparable across AD groups and controls (29-40%) and minimal in FTLD-TDP (8%). Lewy body pathology was present in a subset of AD/LATE cases (17-30%) and infrequently in FTLD-TDP (8-10%) (**Table 1**).

**Table 1.**
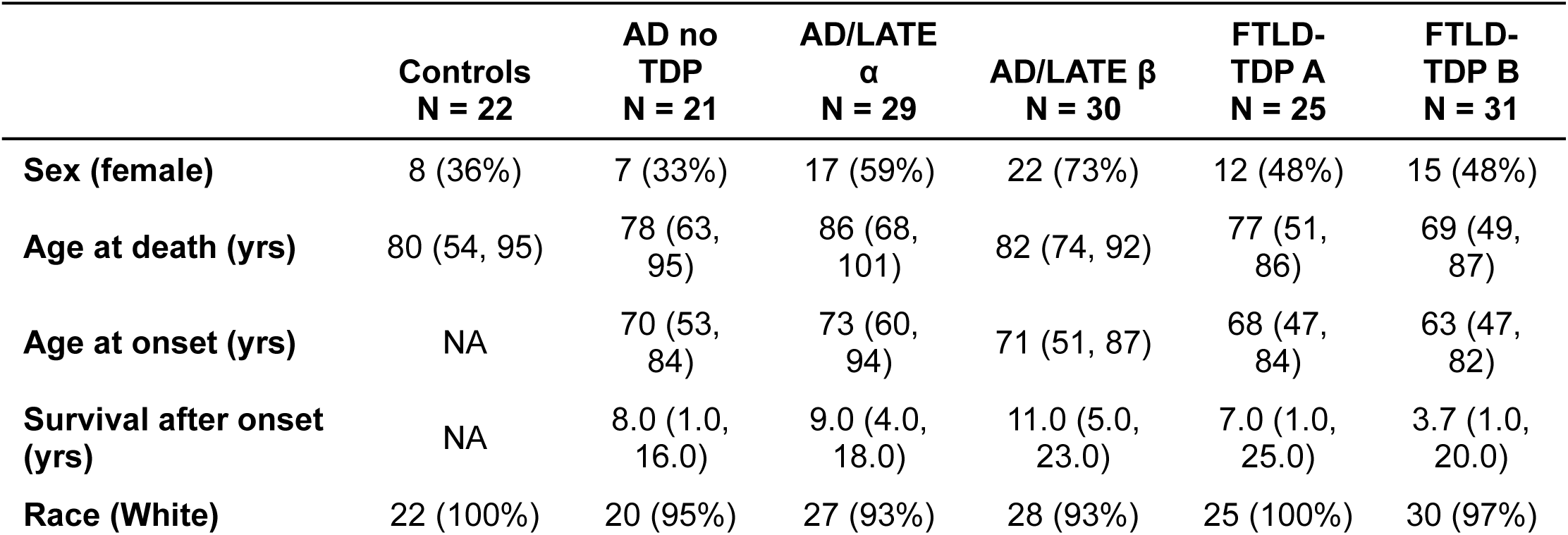
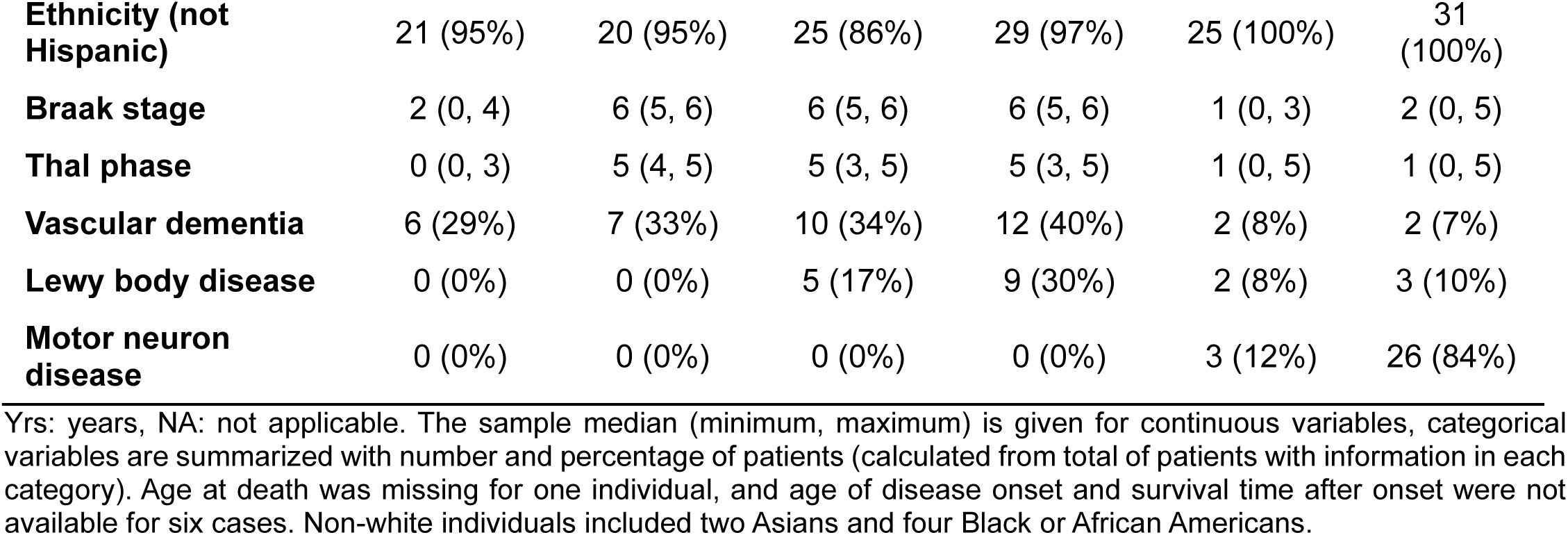
Study cohort characteristics.

| | Controls<br>N = 22 | AD no<br>TDP<br>N = 21 | AD/LATE<br>$\alpha$<br>N = 29 | AD/LATE $\beta$<br>N = 30 | FTLD-TDP A<br>N = 25 | FTLD-TDP B<br>N = 31 |
| --- | --- | --- | --- | --- | --- | --- |
| **Sex (female)** | 8 (36%) | 7 (33%) | 17 (59%) | 22 (73%) | 12 (48%) | 15 (48%) |
| **Age at death (yrs)** | 80 (54, 95) | 78 (63, 95) | 86 (68, 101) | 82 (74, 92) | 77 (51, 86) | 69 (49, 87) |
| **Age at onset (yrs)** | NA | 70 (53, 84) | 73 (60, 94) | 71 (51, 87) | 68 (47, 84) | 63 (47, 82) |
| **Survival after onset (yrs)** | NA | 8.0 (1.0, 16.0) | 9.0 (4.0, 18.0) | 11.0 (5.0, 23.0) | 7.0 (1.0, 25.0) | 3.7 (1.0, 20.0) |
| **Race (White)** | 22 (100%) | 20 (95%) | 27 (93%) | 28 (93%) | 25 (100%) | 30 (97%) |
| Ethnicity (not Hispanic) | 21 (95%) | 20 (95%) | 25 (86%) | 29 (97%) | 25 (100%) | 31 (100%) |
| Braak stage | 2 (0, 4) | 6 (5, 6) | 6 (5, 6) | 6 (5, 6) | 1 (0, 3) | 2 (0, 5) |
| Thal phase | 0 (0, 3) | 5 (4, 5) | 5 (3, 5) | 5 (3, 5) | 1 (0, 5) | 1 (0, 5) |
| Vascular dementia | 6 (29%) | 7 (33%) | 10 (34%) | 12 (40%) | 2 (8%) | 2 (7%) |
| Lewy body disease | 0 (0%) | 0 (0%) | 5 (17%) | 9 (30%) | 2 (8%) | 3 (10%) |
| Motor neuron disease | 0 (0%) | 0 (0%) | 0 (0%) | 0 (0%) | 3 (12%) | 26 (84%) |

Yrs: years, NA: not applicable. The sample median (minimum, maximum) is given for continuous variables, categorical variables are summarized with number and percentage of patients (calculated from total of patients with information in each category). Age at death was missing for one individual, and age of disease onset and survival time after onset were not available for six cases. Non-white individuals included two Asians and four Black or African Americans.

We focused our analysis on the amygdala and frontal cortex because these regions show distinct patterns of TDP-43 involvement: the amygdala is affected in both AD/LATE and FTLD-TDP, whereas the frontal cortex exhibits substantial pathology only in FTLD-TDP^3,9,13^. Given prior evidence that TDP-43 subtypes are associated with specific risk factors and molecular changes in FTLD-TDP^14^, we further stratified AD/LATE cases by TDP-43 subtype to identify transcriptomic features linked to subtype-specific TDP-43 pathology (**Fig. 1a**, **Table 1**). Consistent with previous reports^4,11,12^, histopathological assessment of pTDP-43 and pTau revealed that pTDP-43 inclusions frequently occurred independently of pTau in AD/LATE type α, although adjacent cells could harbor both pathologies without colocalization (**Fig. 1b**). In contrast, AD/LATE type β showed co-localization of pTDP-43 and pTau positive inclusions, interspersed with neighboring cells containing only pTau inclusions (**Fig. 1c**).

**Fig. 1.**
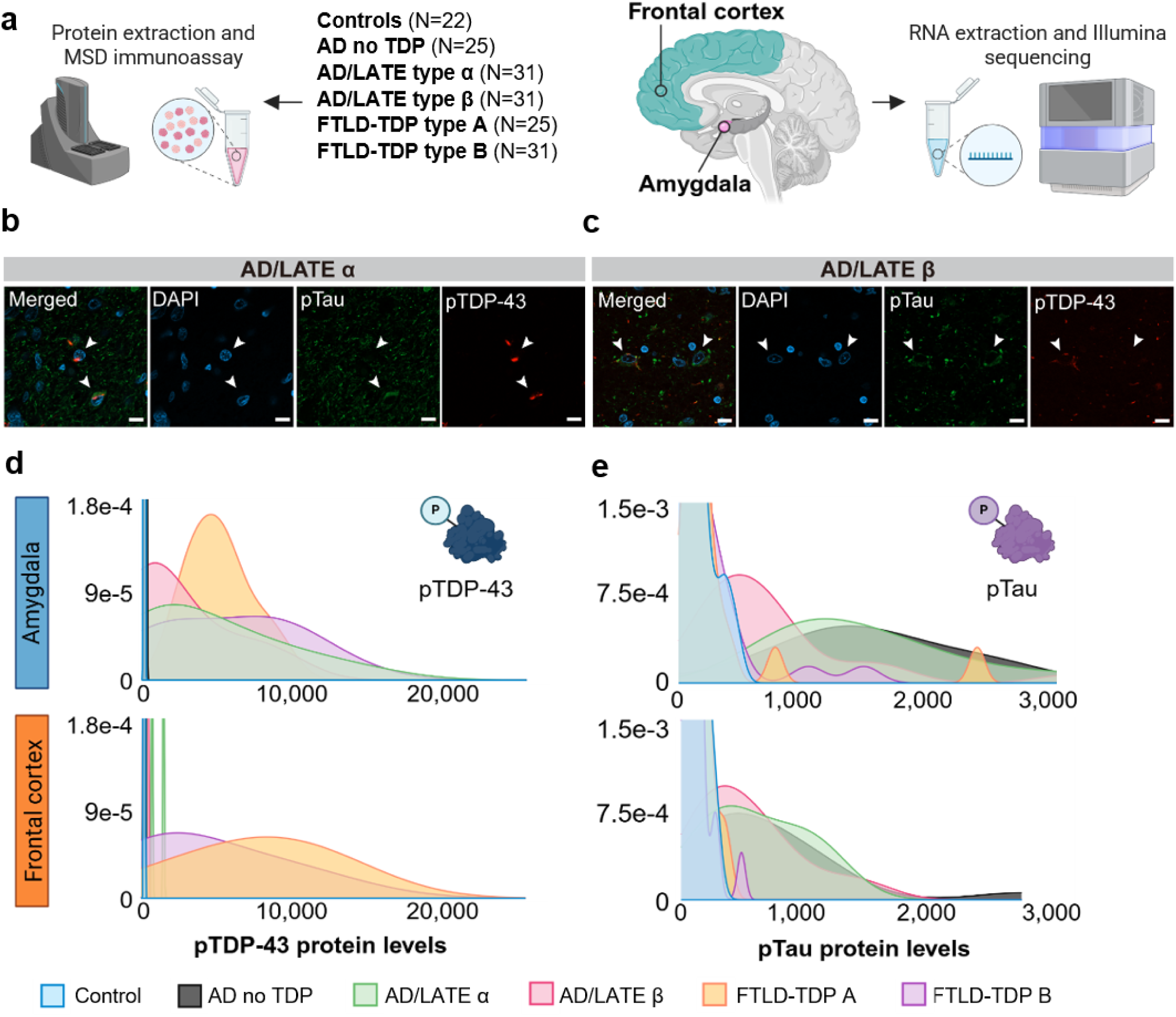
Experimental workflow and quantitative analysis of neuropathological proteins in a human post-mortem cohort. (**a**) Schematic of study design illustrates amygdala and frontal cortex tissues from six neuropathologically confirmed groups. Protein and RNA were extracted to quantify pTDP-43 and pTau protein levels, and to perform RNA sequencing, respectively. (**b, c**) Representative immunofluorescence confocal microscopy images of post-mortem amygdala co-stained for pTau (green), pTDP-43 (red), and nuclei (blue) in AD/LATE type α (**b**) and AD/LATE type β (**c**). Arrows indicate a cell exhibiting independent pTDP-43 pathology, alongside an adjacent cell harboring both pTDP-43 and pTau without intracellular co-localization in AD/LATE type α (**b**). In AD/LATE type β (**c**), arrows highlight a cell displaying distinct intracellular co-localization of fibrillary pTDP-43 and pTau inclusions, observed alongside neighboring cells burdened solely by pTau. (**d, e**) Density distribution plots showing the quantification of pathological pTDP-43 (**d**) and pTau (**e**) proteins, across study groups in the amygdala (top) and frontal cortex (bottom).

Quantification of pathological protein burden further highlighted these distinctions. In the amygdala, pTDP-43 levels were more widespread in AD/LATE type α (median: 2,579; range: 175-15,602) and FTLD-TDP type B (median: 6,684; range: 262-13,924) compared to AD/LATE type β (median: 764; range:156-9,519) or FTLD-TDP type A (median: 4,894; range: 2,325-9,141) (**Fig. 1d, Extended Data Table 1**). In the frontal cortex, pTDP-43 levels were negligible across all AD groups (**Fig. 1d, Extended Data Table 1**), whereas FTLD-TDP cases showed substantial pTDP-43 pathology, with similar ranges between type A (range: 193-14,167) and type B (range: 189-14,622), though type A exhibited higher median levels (7,813 vs. 2,646) (**Fig. 1d, Extended Data Table 1**).

The levels of pTau were high across all AD groups in both brain regions relative to controls and FTLD-TDP cases (**Fig. 1e, Extended Data Table 1**). In the amygdala, pTau burden was highest in AD no TDP (median: 1,477, range: 205-4,452) and AD/LATE type α (median: 1,337, range: 822-3,404), and lower in AD/LATE type β (median: 618, range: 258-3,115). In the frontal cortex, pTau levels were comparable for all three AD groups and similar to those observed in AD/LATE type β amygdala (**Fig. 1e, Extended Data Table1**).

### Transcriptomic signatures define disease and TDP-43 subtypes in AD and FTLD

To examine how TDP-43 dysfunction impacts the molecular signature in AD that may differ from that in controls, AD no TDP, FTLD-TDP, and with respect to TDP-43 subtypes, we first assessed overall transcriptomic similarity using Euclidean distance-based hierarchical clustering from the RNAseq data of the amygdala (**Fig. 2a**) and frontal cortex (**Fig. 2b**). In the amygdala, all AD groups separated from controls, with AD/LATE type α and type β being more similar to each other than to the AD no TDP group (**Fig. 2a**). FTLD-TDP type A and type B in the amygdala also clustered separately from controls and AD/LATE, with FTLD-TDP type B showing more similarity to the AD no TDP group (**Fig. 2a**). In the frontal cortex, where TDP-43 pathology is absent for AD, all AD subtypes clustered more closely, although separated from controls (**Fig. 2b**). FTLD-TDP type A cases in the frontal cortex aligned more closely with AD cases, while FTLD-TDP type B cases clustered closer to controls (**Fig. 2b**).

**Fig. 2.**
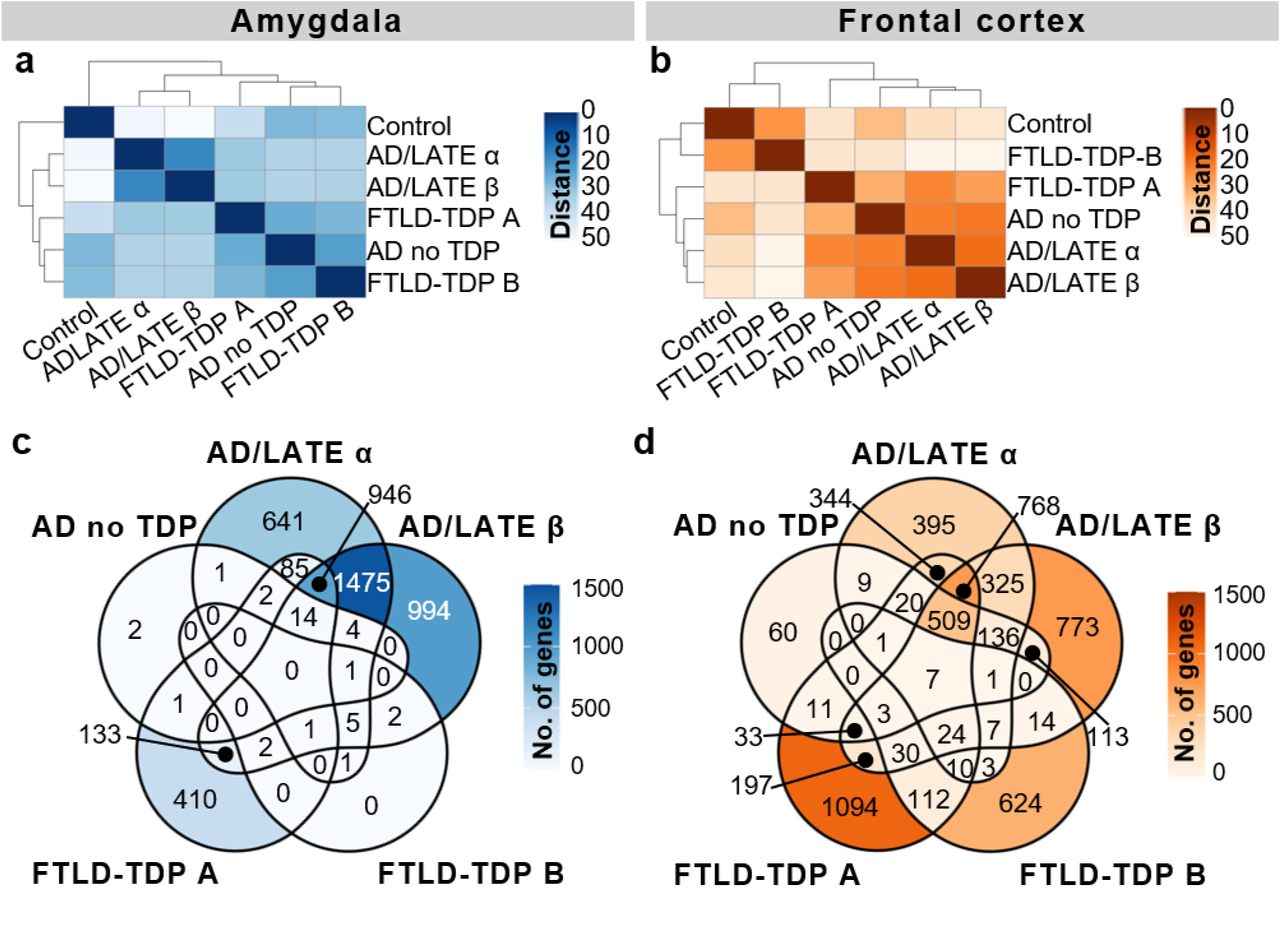
Gene expression changes are regionally and disease-specifically distributed. **(a-b)** Hierarchical clustering heatmaps showing pairwise Euclidian distances between groups, generated from the variance-stabilizing-transformed (VST) expression matrix, for the amygdala (**a**) and frontal cortex (**b**). Color intensity reflects the Euclidian distance between sample pairs, where darker shades indicate lower distances and greater similarity, and lighter shades indicate higher distances and lower similarity. Dendrograms indicate the hierarchical relationship among groups based on transcriptomic similarity. (**c-d**) Venn diagrams depicting the overlap of DEGs across AD/LATE α, AD/LATE β, AD no TDP, and FTLD-TDP subtypes in the amygdala (**c**) and frontal cortex (**d**). The color intensity is reflective of gene number, with darker shades representing groups with greater numbers (No.) of DEGs.

To identify molecular drivers underlying the clustering differences across brain regions and disease subtypes, we next performed differential expression analysis. This analysis revealed 18,785 differentially expressed genes (DEGs) that were significantly dysregulated (adjusted *p*-value < 0.05) by at least 0.5-fold across all disease groups and brain regions (**Supplementary File 1**). The majority of DEGs (N=15,949; 85%) corresponded to protein-coding transcripts, while the rest primarily consisted of long intergenic non-coding RNAs (lincRNAs), antisense RNAs, and various classes of pseudogenes. The total number of DEGs were similarly distributed in the amygdala (N=8,384; 44.6%) and frontal cortex (N=10,401; 55.4%). Within the amygdala, the majority of DEGs resulted from AD/LATE, similarly distributed between type α (N=3,176; 37.9%) and β (N=3,577; 42.7%), and FTLD-TDP type A (N=1,594; 19.0%; **Fig. 2c**). Further, most AD/LATE type α DEG (N=2,421, 76.2%) were shared with those in AD/LATE type β, compared to only 32.5% of DEGs shared with FTLD-TDP type A (**Fig. 2c**). Each subtype also exhibited changes unique to each subtype: 641 (20.2%) for AD/LATE type α, 994 (27.8%) for AD/LATE type β, and 410 (25.7%) for FTLD-TDP type A (**Fig. 2c**). Of interest, the number of DEGs identified in the amygdala of AD no TDP (N=25; 0.3%) and FTLD-TDP type B (N=12; 0.1%) was minimal (**Fig. 2c**). In the frontal cortex, transcriptome changes were identified across all study groups, with lower proportion of changes in AD no TDP (N=903; 8.7%) and FTLD-TDP type B (N=836, 8%) compared to AD/LATE type α (N=2,559; 24.6%), AD/LATE type β (N=2,940; 28.3%), and FTLD-TDP type A (N=3,162; 30.4%) (**Fig. 2d**). The proportion of unique DEG in FTLD-TDP type A (N=1,094; 34.6%) as well as in AD/LATE (α: N=395; 15.4%; and β: N=773; 26.3%) in the frontal cortex (**Fig. 2d**) was similar to the proportions found in the amygdala (**Fig. 2c**).

Across all study groups, there was a larger proportion of downregulated DEGs compared to upregulated DEGs in both the amygdala (**Extended Data Fig. 1a-e**) and frontal cortex (**Extended Data Fig. 1f-j)**. However, this pattern shifted when isolating DEGs unique to each condition. In the amygdala, the proportion of up and downregulated DEGs was relatively balanced for AD/LATE type α (downregulated: N=320, 10.1%; upregulated: N=321, 10.1%), AD/LATE type β (downregulated: N=543, 15.2%; upregulated: N=453, 12.7%) and FTLD-TDP type A (downregulated: N=174, 10.9%; upregulated: N=236, 14.8%), as well as in the frontal cortex for AD no TDP cases (downregulated: N=28, 3.1%; upregulated: N=32, 3.5%) (**Extended Data Fig. 1a-e**). In contrast, the proportions of DEGs unique to each condition varied in the frontal cortex. A higher proportion of downregulated unique DEGs was observed in AD/LATE type α (downregulated: N=265, 10.4%; upregulated: N=130, 5.1%) and FTLD-TDP type B (downregulated: N=498, 53.8%; upregulated: N=126, 13.6%). Conversely, the proportion of unique DEGs shifted in AD/LATE type β (downregulated: N=262, 9.1%; upregulated: N=511, 17.7%) and FTLD-TDP type A (downregulated: N=226, 7.1%; upregulated: N=868, 27.4%), where upregulated unique DEGs constituted a larger proportion of the total (**Extended Data Fig. 1f-j**).

### Gene enrichment analyses of condition-specific DEGs demonstrate shared and unique mechanisms across TDP-43 proteinopathies and subtypes

Gene enrichment analysis of global DEGs across each tissue and condition revealed altered pathways, which were grouped into seven major biological categories: immune and infection, metabolism, gene processing, structural organization, cell processing and development, signaling and neuronal functions (**Extended Data Fig. 2, Supplementary File 2**). To determine molecular pathways specific to each condition, we performed enrichment analysis in condition-specific DEGs across both brain regions (**Fig. 3a**, **Supplementary File 3**). FTLD-TDP type B and AD no TDP were distinguished by broad pathway downregulation in the frontal cortex (**Fig. 3a, Supplementary File 3**), including minimal engagement of immune and infection-related pathways. In contrast, AD/LATE type α, AD/LATE type β, and FTLD-TDP type A showed a distinct and consistent upregulation of immune and infection pathways (**Fig. 3a, Supplementary File 3**). Despite this shared upregulation trend, the specific immune and infection pathways diverged substantially across disease types (**Fig. 3b**). Specifically, AD/LATE type α was defined by an IFN-β-associated antiviral program together with Th2, IL-4, and IL-27 signaling. In contrast, AD/LATE type β engaged an IFN-α-associated innate immune signature, IL-18 signaling, an Fc-receptor-related regulatory module, and NF-κB signaling (**Fig. 3b, Supplementary File 4**). The immune profile of FTLD-TDP type A differed from both AD/LATE subtypes and exhibited regional polarity. In the amygdala, FTLD-TDP type A favored adaptive immune programs, including Th17 cell differentiation and MHC class I biosynthesis (**Fig. 3b, Supplementary File 4**). Conversely, the frontal cortex was dominated by an acute inflammatory response driven by granulocyte and neutrophil activation, degranulation, and migration pathways (**Fig. 3b, Supplementary File 4**).

**Fig. 3.**
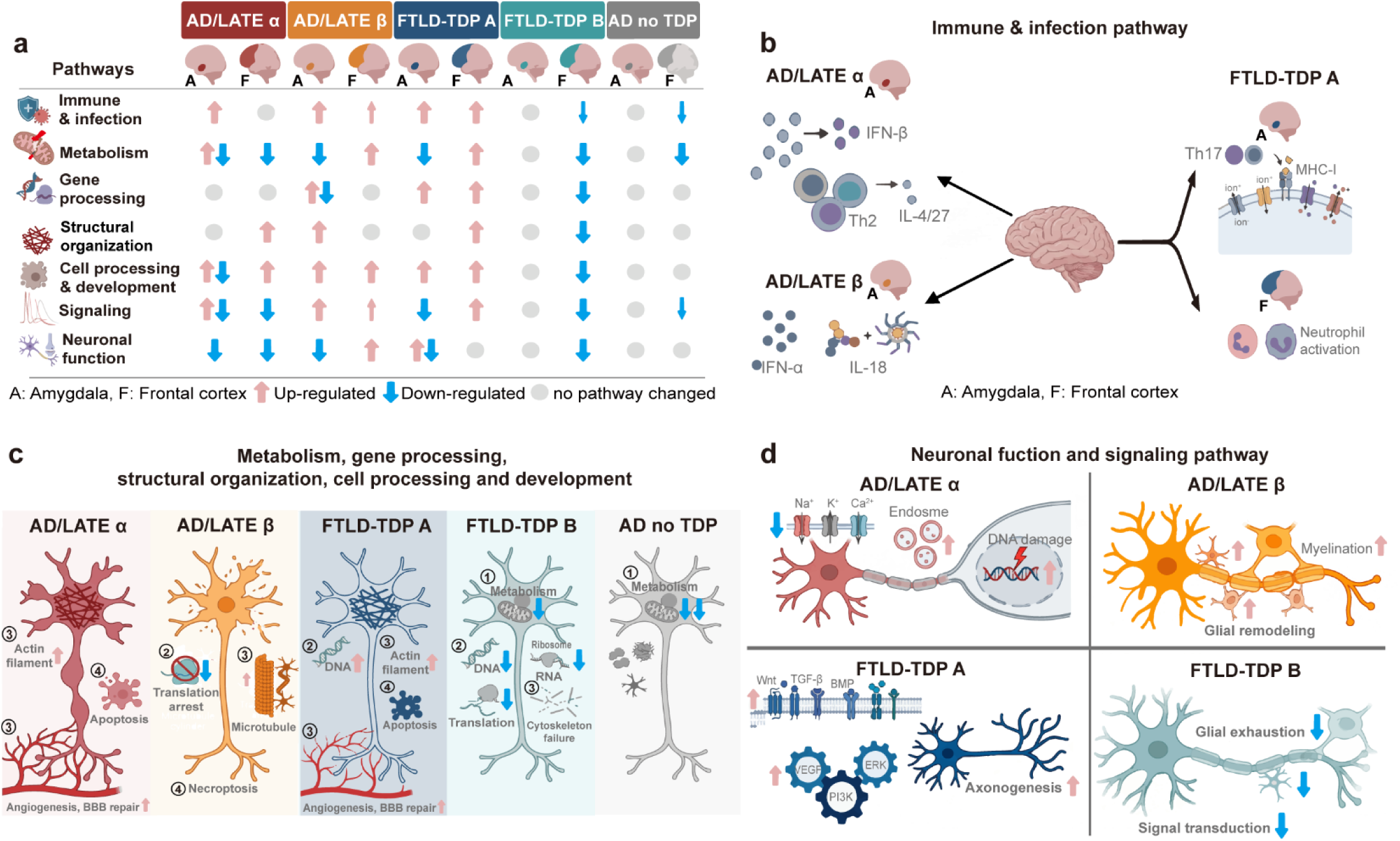
Shared and unique biological mechanisms across TDP-43 proteinopathies and pathological subtypes. (**a**) Global picture of pathway-level perturbations in condition-specific DEGs. Pink upward and blue downward arrows indicate the directional up- and down-regulation of core biological modules, respectively, across the amygdala (A) and frontal cortex (F). Thinner arrows denote categories where only one or two related pathways are altered. Grey dots indicate that no significant pathway changes were detected in that category. (**b**) Condition-specific DEGs related to immune and infection signatures differ among AD/LATE type α, AD/LATE type β and FTLD-TDP type A. (**c**) Condition-specific alterations related to metabolism (1), gene processing (2), structural organization (3), and cell death pathways (4) differ among the disease subtypes. (**d**) Condition-specific disruptions to neuronal function and major signaling cascades displayed distinct, divergent changes across the disease subtypes. AD no TDP is omitted from this panel due to minimal alterations in these pathways.

Across the remaining six pathway categories, AD/LATE type α, AD/LATE type β, and FTLD-TDP type A exhibited a mixture of upregulated and downregulated processes, while FTLD-TDP type B and AD no TDP were again characterized by pathway downregulation in the frontal cortex (**Fig. 3a, Supplementary File 3**). Beyond immune and infection pathways, additional molecular domains further differentiated the disease groups, revealing mechanistically distinct alterations across metabolism, gene regulation, structural organization and cell processing (**Fig. 3c, Supplementary File 4**). Metabolic pathways showed some striking contrast. AD no TDP was defined almost entirely by broad metabolic decline, with reduced activity in amino acid biosynthesis, carbon metabolism, glycolysis/gluconeogenesis, the TCA cycle, NADPH regeneration, and organic acid metabolism (**Fig. 3c, Supplementary File 4**). FTLD-TDP type B exhibited a similarly global metabolic suppression with no concurrent up-regulation of other metabolic pathways (**Fig. 3c, Supplementary File 4**). Conversely, AD/LATE type α, AD/LATE type β and FTLD-TDP type A displayed upregulation of some macromolecule and nucleotide biosynthesis pathways, suggesting an attempt to sustain anabolic processes despite the bioenergetic downregulation within the groups (**Supplementary File 4**).

Differences in gene processing also contributed to the molecular divergence among groups. While no significant pathway alterations were observed in AD no TDP and AD/LATE type α, FTLD-TDP type A exhibited prominent upregulation of transcriptional and nucleic acid-related programs in both brain regions and AD/LATE type β exhibited downregulation of translation only in the amygdala (**Fig. 3c, Supplementary File 4**). The global suppression observed in FTLD-TDP type B frontal cortex was characterized by the concurrent downregulation of RNA processing, ribosome assembly, translation, and ubiquitin-mediated protein regulation pathways (**Fig. 3c, Supplementary File 4**).

Structural organization pathways provided another axis of separation. AD/LATE type α and FTLD-TDP type A both upregulated actin cytoskeleton remodeling, cell adhesion, and cellular motility programs, which was further characterized by a shared hyper-reactive neurovascular remodeling trajectory (**Fig. 3c, Supplementary File 4**). In contrast, FTLD-TDP type B broadly downregulated pathways involved in cellular component assembly and organization (**Fig. 3c, Supplementary File 4**). AD/LATE type β stood apart by uniquely upregulating microtubule- and cilia-related pathways, while AD no TDP lacked changes in this pathway category (**Fig. 3c, Supplementary File 4**).

Within cell processing and development categories, cell death mechanisms mainly diverged between AD/LATE subtypes. AD/LATE type α was associated with classical apoptotic pathways, which was shared with FTLD-TDP type A, whereas AD/LATE type β engaged necroptotic cell death (**Fig. 3c, Supplementary File 4**), underscoring fundamentally different modes of neurodegeneration within the LATE spectrum.

Finally, neuronal function and signaling pathways provided yet another layer of subtype-specific remodeling, extending the molecular distinctions observed across immune, metabolic, and regulatory domains (**Fig. 3d, Supplementary File 4**). Both AD/LATE type α in the amygdala and FTLD-TDP type A in the frontal cortex shared an upregulation of genes involved in MAPK and integrin signaling (**Supplementary File 4**). However, the FTLD-TDP type A showed a further upregulation of canonical Wnt, BMP, TGF-β, VEGF, PI3K, and ERK cascades occurring alongside markers of axonogenesis in the frontal cortex (**Fig. 3d, Supplementary File 4**). Despite these shared signaling activations, both AD/LATE type α and FTLD-TDP type A displayed a suppression of core ion-handling functions across brain regions. However, AD/LATE type α uniquely exhibited a concurrent upregulation of ion homeostasis, endocytosis, and positive transport regulatory programs, as well as DNA damage-responsive signaling and integrity checkpoints (**Fig. 3d, Supplementary File 4**). Furthermore, AD/LATE type β again stood apart, showing enrichment for pathways associated with collateral sprouting, radial glial scaffold formation, and central nervous system myelination. In contrast, FTLD-TDP type B was defined by broad downregulation of gliogenesis, glial cell differentiation, signal transduction cascades, and oligodendrocyte development (**Fig. 3d, Supplementary File 4**).

### TDP-43 dysfunction drives distinct transcriptomic changes in AD/LATE independent of pTau

To disentangle the specific molecular contributions of TDP-43 from co-occurring Tau pathology within these AD/LATE subtypes, we integrated the transcriptomic data with quantitative biochemical measures of pTDP-43 and pTau in the amygdala by performing biweight midcorrelation (BiCor) analysis. Notably, the majority of pTDP-43-correlated DEGs in AD/LATE type α (N=223; 59.3%) and 70.7% in AD/LATE type β (N=263; 70.7%), did not correlate with pTau (**Fig. 4a**). Conversely, 78.6% of pTau-correlated DEGs (N=562) in AD/LATE type α and 82.6% in AD/LATE type β (N=523) did not correlate with pTDP-43 (**Fig. 4a-b**).

**Fig. 4.**
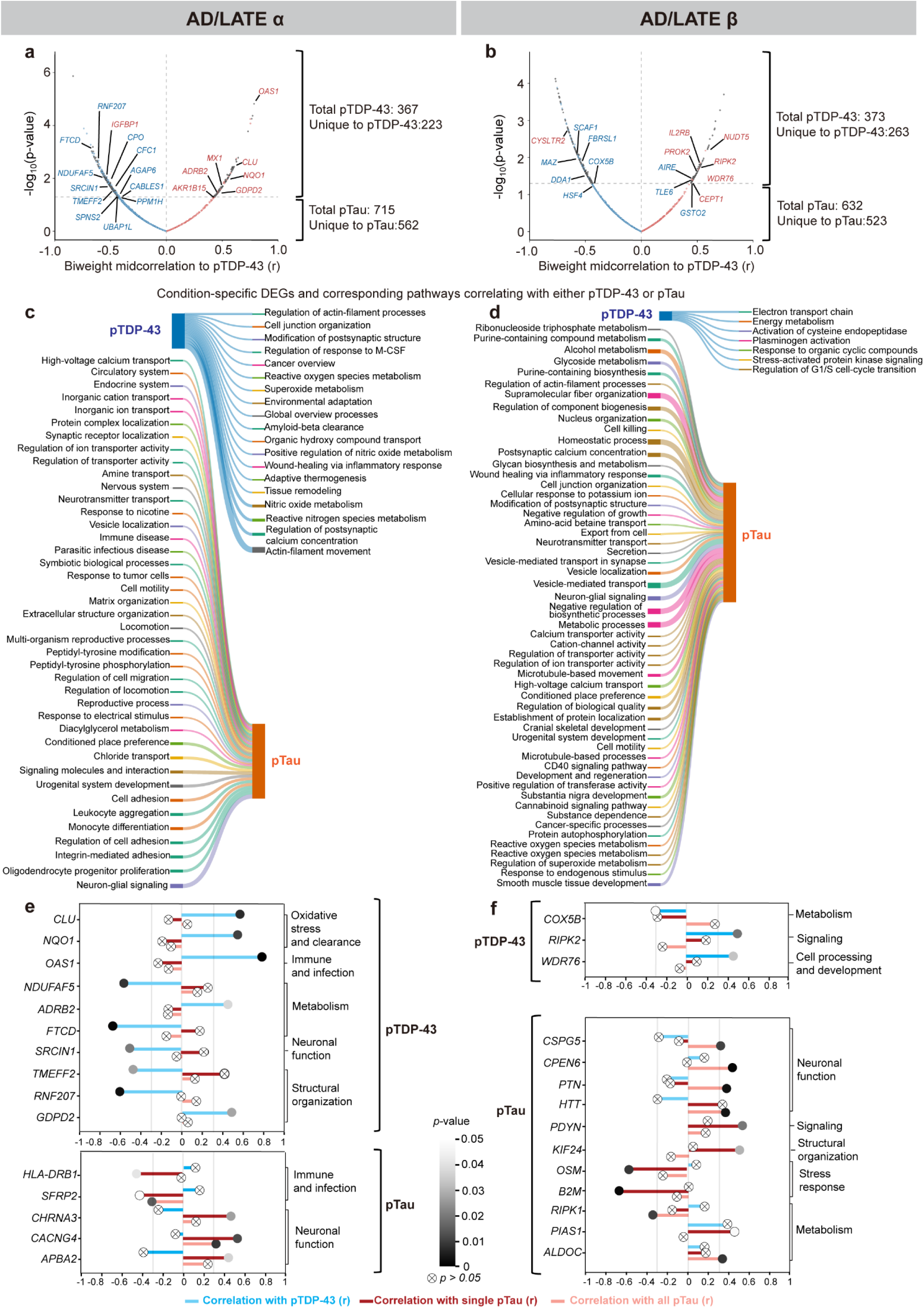
Subtype-specific pathway enrichment and transcriptomic uncoupling of pTDP-43 and pTau pathologies in the AD/LATE amygdala. (**a, b**) BiCor volcano plots showing the strength and significance of the correlation between DEG expression and pTDP-43 levels in (**a**) AD/LATE α (20 annotated DEGs labeled) and (**b**) β (16 annotated DEGs labeled). The x-axis represents the biweight midcorrelation coefficient (*r*) and the y-axis represents -log_10_(*p*-value). DEGs represented by grey dots with black outlines are exclusively correlated with pTDP-43 and not pTau; among these, the labeled DEGs are unique to the condition and the amygdala. Gene names in blue indicate downregulation; red indicates upregulation. Labeled DEGs are significantly correlated |r| > 0.3, adjusted *p*-value<0.05). (**c, d**) Sankey diagrams illustrating pathway-level associations uniquely driven by pTDP-43 (left nodes) or pTau (right nodes) in the AD/LATE type α (**c**) and type β (**d**) amygdala. Uniquely correlated gene sets were mapped to subtype-enriched pathways and consolidated via REVIGO (detailed in **Supplementary File 5**). Flow thickness represents the relative contribution of genes supporting each biological module. (**e–f**) Correlation coefficient plots visualizing the specific BiCor values (r) for representative pathway-defining genes in AD/LATE type α (**e**) and type β (**f**). The x-axis denotes the correlation coefficient (r). Colored lines and nodes denote the specific correlation tested: cyan indicates correlation with pTDP-43; dark red indicates correlation with subtype-specific pTau (single pTau); light red indicates correlation with overall pTau in all AD cases (all pTau). Node size and grayscale fill intensity represent statistical significance (adjusted *p*-value), with crossed-out circles denoting non-significant associations (*p*-value > 0.05).

Since TDP-43 dysfunction in AD is restricted to the amygdala, we further refined our analysis by focusing on condition-specific DEGs that are uniquely dysregulated in the amygdala, such that they are not dysregulated in the same direction in any other condition or brain region. This stringent filtering revealed 31 unique DEGs in AD/LATE type α (labeled in **Fig. 4a**) and 17 unique DEGs in AD/LATE type β (labeled **Fig. 4b**) that correlate exclusively with pTDP-43 burden (**Fig. 4a-b, Supplementary File 5**). This approach also revealed 26 pTau-exclusive DEGs (not correlated with pTDP-43) in the AD/LATE type α amygdala (labeled in **Extended Data Fig. 3a-b**) and 79 in the AD/LATE type β amygdala (labeled in **Extended Data Fig. 3c-d**), however, these were not filtered against frontal cortex expression, as Tau pathology is characteristically present in that region in AD (**Fig. 4a-b, Supplementary File 5**).

We then determined which molecular pathways are associated with condition-specific pTDP-43 or pTau correlated DEGs. We found that uniquely correlated genes often converged on shared biological networks, broadly encompassing general metabolism, structural remodeling, and inflammatory responses in both subtypes (**Extended Data Fig. 4a-d**). However, each pathology also displayed distinct, unique pathway-level associations (**Fig. 4c-d**). For example, in AD/LATE type α, pTDP-43 uniquely associated with pathways related to amyloid-beta clearance, reactive oxygen and nitrogen species metabolism, and wound healing (**Fig. 4c**) while, in AD/LATE type β, pTDP-43 was associated with energy metabolism and the activation of endopeptidase activity (**Fig. 4d**). Conversely, pTau was associated with numerous uniquely misregulated pathways (**Fig. 4c-d**). In AD/LATE type α, pTau was linked to high-voltage calcium transport, neurotransmitter transport, and neuron-glial signaling (**Fig. 4c**), while the AD/LATE β subtype associated with a wide array of structural and regulatory pathways, including microtubule-based movement, vesicle-mediated transport, and homeostatic processes (**Fig. 4d**).

Next, we sought to identify the specific genes driving these distinct pathology-exclusive mechanisms. In AD/LATE type α, pTDP-43-associated increased activity in stress- and immune-related pathways was linked to the upregulation of several major AD risk genes, including *CLU, NQO1,* and *OAS1* (**Fig. 4e**). pTDP-43-correlated DEGs associated with reduced activity in pathways related to environmental adaptation, cellular metabolism and tissue remodeling included *NDUFAF5*, *FTCD* and *ADRB2* (**Fig. 4e**). Reduced activity in pathways involved in postsynaptic structural regulation was associated with the downregulation of *SRCIN1*, whereas decreased activity in pathways related to actin filament organization and cell adhesion was associated with *TMEM2*, *RNF207*, and *GDPD2* (**Fig. 4e**).

In AD/LATE type β, pTDP-43-correlated DEGs associated with reduced activity in mitochondrial energy metabolism pathways included *COX5B* (**Fig. 4f**). In contrast, increased activity in stress-responsive signaling pathways was associated with *RIPK2*, while enhanced activity in pathways related to G1/S cell-cycle transition and DNA damage responses was associated with *WDR76* (**Fig. 4f**).

There were more condition-specific pathways associated with pTau-related DEGs in AD/LATE than with pTDP-43-associated alterations (**Fig. 4e-f**). In addition, transcriptional changes in AD/LATE type α were relatively limited compared to those observed in type β (**Fig. 4e-f**). In AD/LATE type α, higher pTau levels were associated with increased expression of *HLA-DRB1,* a known AD risk gene, and *SFRP2*, both of which were linked to immune-related pathways (**Fig. 4e**). Further, pTau positively correlated with genes associated with the downregulation of synaptic transport and neuron-glial signaling networks (*CACNG4, CHRNA3, APBA2*) (**Fig. 4e**). In contrast to AD/LATE type α, structural, neuronal, and stress dysregulations in the AD/LATE β amygdala were profoundly correlated with higher pTau levels (**Fig. 4f**). This included the suppression of essential signaling mediators, calcium transport, and postsynaptic structures (*CPNE6*, *CSPG5*, *HTT*, *PTN*, *PDYN*), upregulation of genes such as *KIF24* linked to the altered structural organization and microtubule dynamics, the upregulation of distinct inflammatory and cell-killing regulators (*B2M* ,*OSM*, *RIPK1)* as well as genes associated with disruption of cellular metabolism (*ALDOC, PIAS1*) (**Fig. 4f**).

## Discussion

TDP-43 proteinopathy shows distinct pathological subtypes, neuroanatomical distributions, and clinical manifestations across neurodegenerative diseases. In AD/LATE, TDP-43 pathology is confined to limbic regions, whereas in FTLD-TDP it extends more broadly into neocortical areas, including the frontal cortex. These differences in anatomical distribution have been associated with differences in clinical presentation and disease progression^3,9^. By comparing AD/LATE and FTLD-TDP cases across brain regions and TDP-43 subtypes, we found that while transcriptional profiles retain distinct disease-specific features, the magnitude of regional transcriptomic remodeling is shaped by local pTDP-43 burden and subtype. In the amygdala, substantial transcriptomic alterations were observed in both AD/LATE subtypes and in FTLD-TDP type A, all of which showed prominent local pTDP-43 pathology. In contrast, AD cases lacking TDP-43 pathology and FTLD-TDP type B cases showed relatively few transcriptional changes in this region. A different pattern emerged in the frontal cortex, where transcriptional alterations were most pronounced in FTLD-TDP, particularly type A, consistent with its high neocortical pTDP-43 burden. Although frontal cortex changes were also detected in AD/LATE, pTDP-43 pathology was largely absent from this region, suggesting that these alterations are more likely attributable to AD-related processes than to local TDP-43 pathology. Together, these findings indicate that the molecular consequences of TDP-43 pathology are closely linked to its regional distribution. By distinguishing TDP-43-associated transcriptional signatures from the effects of coexisting tau pathology, our results suggest that TDP-43 is not simply an accompanying pathology in AD. Instead, it is associated with distinct molecular changes that vary across brain regions and pathological subtypes, potentially contributing to the biological heterogeneity of AD/LATE.

Large-scale transcriptomic and single-nucleus RNA sequencing studies have consistently identified increased neuroinflammatory signaling together with reduced synaptic and mitochondrial function as core molecular features of the AD brain^15–17^. Because AD has traditionally been defined by amyloid and tau pathology, these transcriptional changes have largely been interpreted as downstream consequences of these hallmark lesions. Our findings suggest a more nuanced view. By stratifying AD/LATE cases according to TDP-43 morphological subtype and separating the transcriptional effects associated with pTau and pTDP-43 burden, we found that several molecular signatures commonly regarded as characteristic of AD are also influenced by TDP-43 pathology.

This was particularly evident for immune-related pathways. Neuroinflammation is a well-established feature of AD, and previous transcriptomic studies have identified disease-associated microglial (DAM) states^18^ and interferon-responsive programs as prominent components of the late-onset AD immune response^19,20^. While our data are consistent with these observations, they further suggest that the nature of the inflammatory response varies according to TDP-43 subtype. In the AD/LATE type α amygdala, immune-related changes were characterized by activation of IFN-β-associated antiviral pathways, including the induction of genes such as *OAS1* and *MX1*, together with enrichment of Th2/IL-4 signaling. Notably, these signatures were more strongly associated with pTDP-43 than with pTau burden. The prominence of interferon-responsive pathways is consistent with experimental studies showing that TDP-43 dysfunction can trigger innate immune activation through mechanisms such as cGAS–STING signaling^21^. In parallel, the enrichment of Th2/IL-4-related pathways may reflect a more immunoregulatory microglial response, potentially related to tissue repair or adaptation to early cellular stress^22,23^.

In contrast, the AD/LATE type β subtype showed enrichment of IFN-α, IL-18, and NF-κB signaling pathways, indicating a distinct inflammatory profile. Although transcriptomic data cannot directly assess inflammasome activation, the prominent IL-18 signature is consistent with inflammasome-associated inflammatory responses and aligns with experimental studies linking innate immunity to tau pathology^24^. Notably, despite lower median amygdala pTau levels than other AD groups, type β showed widespread transcriptional alterations that were strongly associated with pTau burden. Together with the activation of NF-κB, a master regulator of chronic inflammation^25^, these findings suggest a more pro-inflammatory immune state in type β compared with type α.

Beyond the interferon-driven responses observed in AD/LATE, FTLD-TDP type A shows a distinct, adaptive immune profile. In the FTLD-TDP type A amygdala, the immune profile features adaptive immune responses, particularly Th17 signaling and MHC-I upregulation. The increase in MHC-I suggests enhanced antigen presentation and immune surveillance, while the associated Th17 response is a recognized pathway implicated in neuroinflammation^26^. Conversely, the frontal cortex in FTLD-TDP type A shows enrichment of pathways related to neutrophil activation and degranulation^27^. Because neutrophils are acute responders that do not typically reside in a healthy brain, the upregulation of these transcripts is consistent with innate immune activation and localized tissue stress. Collectively, these findings indicate that TDP-43 pathology is associated with distinct, subtype-specific inflammatory programs rather than a single generalized neuroinflammatory response.

We also identified subtype-specific patterns of neuronal and mitochondrial dysfunction, suggesting that TDP-43 and tau pathologies affect distinct biological processes within the brain. Although AD no TDP cases already showed evidence of impaired energy metabolism, the addition of TDP-43 was associated with more clearly differentiated mechanisms of neurodegeneration. In the AD/LATE type α amygdala, the effects of pTDP-43 and pTau appeared largely segregated. pTDP-43 was primarily associated with the loss of transcripts involved in cellular maintenance and neuroprotection, whereas pTau was more closely linked to deficits in neuronal excitability and cholinergic signaling. By contrast, the AD/LATE type β subtype exhibited a more coordinated pattern in which pTau emerged as the major driver of synaptic dysfunction and cytoskeletal disruption. In this context, pTau was associated with broad suppression of synaptic mediators and trophic support pathways, while pTDP-43 contributed a more selective vulnerability by targeting genes involved in oxidative phosphorylation and mitochondrial energy production. This metabolic signature aligns with experimental evidence implicating TDP-43 dysfunction in mitochondrial impairment across the ALS and FTLD spectrum^28,29^.

Crucially, the distinct immune and metabolic pathways include genes previously implicated in late-onset AD risk, suggesting potential links between genetic susceptibility and subtype-associated molecular programs. In AD/LATE type α, the pTDP-43-associated oxidative stress and antiviral response pathways involve both established and putative risk genes, including *CLU*, *NQO1*, and *OAS1*. Genetic variants at the *CLU* locus strongly influence AD susceptibility by modulating both protein clearance^30^ and inflammatory responses^31^, with prior evidence suggesting a role for clusterin in clearing TDP-43 aggregates^32^. Similarly, specific polymorphisms in *NQO1* have been associated with altered AD risk in certain populations, though further validation is needed^33–35^. The relationship between pTDP-43 and innate immune activation is further underscored by its unique correlation with *OAS1*, a recognized GWAS risk gene^36^, supporting the clinical relevance of this interferon-associated stress response. Conversely, the pTau-associated pathways within the α subtype selectively engaged *HLA-DRB1*. As a core component of the microglial MHC Class II complex and a recognized genetic risk factor for late-onset AD^37,38^, its exclusive correlation with pTau suggests that adaptive immune modulation and antigen presentation may be transcriptionally distinct from the pTDP-43-associated innate antiviral-like response. In AD/LATE type β, the specific impairment of oxidative phosphorylation pathways associated with pTDP-43 was characterized by the downregulation of *COX5B*. This aligns with the broader oxidative phosphorylation deficits and reduced mitochondrial gene expression widely reported across AD, ALS, and FTLD^39–41^, and notably, variants in *COX5B* itself have been linked to AD susceptibility^42^.

Beyond individual genetic pathways, analyzing the aggregate transcriptomic output reveals a shift in the relative transcriptomic association of pTDP-43 and pTau across subtypes. Despite a lower overall pTDP-43 burden in type β cases, the amygdala in AD/LATE type β exhibited a greater number of unique DEGs and a much higher proportion of transcripts correlated with pTau burden. This transcriptomic shift closely mirrors the distinct histological features of these subtypes. In AD/LATE type α, TDP-43 inclusions often occur independently of tau pathology, correlating with segregated transcriptomic effects. In contrast, the diffuse or fibrillary TDP-43 inclusions in AD/LATE type β closely co-mingle with tau neurofibrillary tangles. This spatial co-occurrence may promote synergistic neurotoxicity, explaining why pTau shows a dominant transcriptomic association over pTDP-43 in AD/LATE type β.

A similar relationship between pathological burden and transcriptional changes was also observed in FTLD-TDP. Despite higher amygdala pTDP-43 burden in FTLD-TDP type B compared to FTLD-TDP type A, type B exhibited markedly fewer differentially expressed genes. This non-linear relationship between pathological burden and transcriptomic result indicates that raw pTDP-43 burden alone does not dictate the magnitude of molecular dysregulation, suggesting that subtype-specific biological context, cellular vulnerability, or pathological composition can modify the transcriptional consequences of TDP-43 pathology. Importantly, these findings align with recent large-scale RNA-sequencing studies of FTLD-TDP, which have similarly demonstrated that FTLD-TDP subtypes harbor inherently distinct molecular signatures. For instance, recent transcriptomic evaluations have highlighted that FTLD-TDP type A possesses a uniquely robust transcriptomic and splicing signature, particularly regarding immune and structural pathways, that clearly differentiates it from type B, type C, and other genetic subtypes^43–45^. The distinct transcriptomic responses elicited by these subtypes may be fundamentally rooted in the underlying structural conformations of the protein aggregates. Indeed, recent cryo-electron microscopy studies have revealed that TDP-43 forms distinct amyloid fibril folds across different neurodegenerative diseases^46^, providing a structural and mechanistic basis for why RNA processing and transcriptomic alterations diverge so significantly across subtypes.

Recent bulk RNA-seq meta-analyses of the AD temporal lobe have identified the upregulation of risk genes like CLU as a generalized disease signature ^47^. Prior transcriptomic studies in AD have often treated TDP-43 pathology as a binary feature and have lacked focus on limbic regions such as the amygdala, one of the earliest and most vulnerable sites of TDP-43 pathology. To address these limitations and resolve these complex, burden-dependent changes, our study utilizes a BiCor framework to relate continuous measures of pTDP-43 and pTau pathology to gene expression. This represents a substantial advancement over prior binary case-control comparisons. Crucially, rather than artificially removing pTau-associated signals, this approach simultaneously evaluates both pathologies, allowing us to accurately classify transcripts into uniquely correlated or jointly overlapping pathways. By using this integrated framework, we demonstrate that pTDP-43- and pTau-associated transcriptional changes are both overlapping and non-overlapping in AD/LATE. Strikingly, our analysis reveals that most pTDP-43-correlated genes were not significantly associated with pTau burden. This finding challenges the prevailing assumption that TDP-43–linked transcriptional changes in AD merely reflect downstream consequences of Tau pathology and instead supports a model in which TDP-43 independently engages immune, metabolic, and neuronal signaling programs in a subtype-dependent fashion.

The identification of these subtype-specific molecular pathways carries implications for clinical translation and the design of future therapeutic interventions. Currently, the classification of TDP-43 subtypes in AD/LATE is restricted to postmortem pathological evaluation, an assessment that is not routinely performed across all clinical settings. However, the distinct transcriptomic signatures uncovered here provide a crucial foundation for developing *in vivo* fluid biomarkers. For instance, the translation products of novel cryptic exons generated by TDP-43 loss-of-function could potentially be isolated from cerebrospinal fluid or blood^48,49^. Translating this postmortem molecular dichotomy into living patient biomarkers would represent an important advancement, enabling the antemortem stratification of the AD spectrum. Furthermore, this molecular uncoupling highlights the urgent need for precision medicine in AD clinical trials. The treatment of unstratified cohorts harboring fundamentally divergent pathogenic drivers may be a contributing factor to variable therapeutic responses in AD trials^50^. Our data suggests that therapeutic interventions may need to be tailored to the specific pathological landscape of the patient. For example, pharmacological inhibitors of necroptosis, such as RIPK1 inhibitors currently in clinical development^51,52^, may be efficacious in combating the unique pTau-associated cell-death cascades that dominate the transcriptomic landscape of the AD/LATE type β. However, these same inhibitors would likely provide no clinical benefit to patients with the type α subtype, where the dominant transcriptomic signature is enriched for interferon-associated antiviral-like pathways rather than necroptosis.

Despite these insights, our study has certain limitations. While single-nucleus RNA-sequencing has been extensively applied to AD cohorts to map cellular vulnerabilities, existing publicly available datasets^17,53^ lack detailed neuropathological classification of TDP-43 presence, let alone specific AD/LATE morphological subtypes. Validating our findings in larger, independent cohorts is a critical next step; however, because current large-scale datasets lack this essential neuropathological metadata, generating new single-cell datasets from highly characterized, pathologically stratified cohorts will be required to delineate the precise roles of neurons, microglia, and astrocytes in driving these subtype-specific programs. Second, while our cohort is neuropathologically well-defined, it predominantly consists of White individuals, highlighting the need for future validation in more diverse populations to ensure the generalizability of these molecular axes across different ancestral backgrounds. Finally, the cross-sectional nature of postmortem transcriptomics allows for the identification of robust associations but limits our ability to definitively establish the temporal causality between the emergence of specific TDP-43 subtypes and the initiation of distinct transcriptomic programs.

Collectively, our results indicate that TDP-43 pathology is associated with distinct but partially overlapping molecular programs across AD/LATE subtypes and FTLD-TDP. Together, these findings indicate that AD/LATE cases represent a molecularly distinct state within the clinicopathologic spectrum. Ultimately, our work supports a model in which TDP-43 pathology is associated with recurring transcriptional features that are modified by regional vulnerability, pathological subtype, and its unique interplay with co-occurring proteinopathies.

## Methods

### Study approval and sample selection

Human postmortem brain tissues from the amygdala and frontal cortex were obtained from the Mayo Clinic Florida Brain Bank. Clinical and neuropathological diagnoses were independently established by trained neurologists and neuropathologists based on standardized neurological assessments and postmortem examinations, respectively. Written informed consent was obtained from all participants or their next of kin. Protocols were approved by the Mayo Clinic Institutional Review Board and Ethics Committee. Tissue samples were selected based on confirmed neuropathological diagnoses, including FTLD-TDP, AD with or without TDP-43 pathology, and cognitively normal controls. The number of cases per group was based on available tissue for each diagnostic category. Both male and female donors were included in the study. Cohort details are summarized in **Table 1**.

### Protein extraction from postmortem brain tissue

Postmortem brain samples were dissected from the amygdala and frontal cortex as previously described^13^. For protein extraction, 50–60 mg of tissue from each region was used. The amygdala was obtained from a coronal section at the level of the uncus, and the frontal cortex was taken from Brodmann area 9 (middle frontal gyrus). Protein was extracted from the dissected tissue using a sequential fractionation protocol as previously described ^13^. Briefly, tissue homogenization was performed in cold RIPA buffer (25 mM Tris-HCl, pH 7.6; 150 mM NaCl; 1% sodium deoxycholate; 1% Nonidet P-40; 0.1% SDS) supplemented with protease and phosphatase inhibitors. Homogenates were sonicated on ice and centrifuged at 100,000 × g for 30 minutes at 4 °C. The supernatant was collected as the RIPA-soluble fraction. The remaining pellet was washed, sonicated again in RIPA buffer, and centrifuged under the same conditions. The pellet was then resuspended in urea buffer (30 mM Tris-HCl, pH 8.5; 7 M urea; 2 M thiourea; 4% CHAPS), incubated for 1 hour at room temperature with agitation, sonicated, and centrifuged again at 100,000 × g for 30 minutes at 22 °C. The final supernatant, representing the urea-soluble (RIPA-insoluble) protein fraction, was collected for downstream analyses. Protein concentrations were measured using the Bradford assay (ThermoFisher).

### Quantification of phosphorylated TDP-43

Phosphorylated TDP-43 (pTDP-43) levels were quantified in the urea-soluble protein fraction from the amygdala and frontal cortex using samples from the cohort previously described in Estades et al. (2023) as previously described using a Meso Scale Discovery (MSD) electrochemiluminescence-based immunoassay^13^. In brief, a rabbit polyclonal antibody targeting pTDP-43 at serines 409/410 (3 µg/mL; Proteintech, 22309-1-AP) was used for capture, and a sulfo-tagged rabbit polyclonal antibody recognizing the C-terminus of TDP-43 (3 µg/mL; Proteintech, 12892-1-AP) was used for detection. Each sample was assayed in duplicate, and internal controls were included on each plate to account for interplate variability. Signal detection was performed using MSD QUICKPLEX SQ120.

### Quantification of phosphorylated Tau

Phosphorylated Tau (pTau) protein levels in the amygdala and frontal cortex were quantified using an MSD-based immunoassay. Briefly, using 0.02 ug of protein from RIPA-soluble fraction, pTau was quantified using a mouse monoclonal antibody specific to Tau phosphorylated at S262 and/or S356 (12E8^54^) as the capture antibody (1:500), and E1 (1:500; in-house antibody against amino acid residues 19–33 within exon 1 of human Tau: GLGDRKDQGGYTMHQ^55^) paired with Sulfo-tagged anti-rabbit HRP (500 μg/mL; Meso Scale Discovery, R32AB-1) was used for detection. All samples were tested in duplicate, and internal controls were included on each plate to account for any interplate variability. MSD QUICKPLEX SQ120 technology was used for signal detection.

### Immunofluorescence staining of postmortem human amygdala

Paraffin-embedded amygdala sections were deparaffinized in xylene and rehydrated in a graded series of alcohols. Following several washes, antigen retrieval was performed by steaming in home-made cis-tris buffer (pH 6.0) for 30 minutes. Slides were then washed in distilled water for 20 minutes to allow for cooling, and subsequently blocked with Protein Block (Agilent Dako, X0909) at room temperature for 1 hour. Sections were then incubated overnight at 4°C with primary antibodies against pTau (AT8) (1:1000, ThermoFisher, MN1020AT8) and pTDP-43 (1:2000, 26H10^56^) diluted in Antibody Diluent (S3022, Agilent Dako). After overnight incubation, the slides were washed with 1x TBST (TBS with 0.01% Triton-X) and incubated in secondary antibody conjugated with Alexa Fluor donkey-anti rabbit 488 (1:500, Invitrogen, A21206) or Alexa Fluor donkey-anti mouse 568 (1:500, Invitrogen, A10037) in Antibody Diluent (S3022, Agilent Dako) for 2 hours at room temperature in the dark. Following incubation with the secondary antibody, slides were washed in TBST and then incubated in Hoechst 33258 (1:1000, ThermoFisher, H3569) for 10 minutes in the dark at room temperature. After washing in TBST, autofluorescence was reduced using Sudan Black dye and the slides were coverslipped with Fluoromount G and the images were obtained and processed on a Zeiss LSM 980 laser scanning confocal microscope.

### Visualizing pTDP-43 and pTau distributions by group

To visualize the distribution of pTDP-43 and pTau neuropathologic protein levels in the amygdala and frontal cortex, we generated kernel density (KDE) plots using the ggplot2^57^ package in R (v. 4.4.1)^58^. For each tissue and measured protein, group-specific density curves for control, AD no TDP, AD/LATE type α, AD/LATE type β, FTLD-TDP type A, and FTLD-TDP type B were overlaid within the same plot, with each density estimate scaled to unit area (area under each curve = 1).

### RNA isolation from human brain samples

40 mg of the RNA was extracted from human brain samples was used for RNA isolation, as previously described^13^. In brief, the RNeasy Plus Mini Kit (Qiagen) was used to extract RNA according to the manufacturer’s instructions. Up to three cuts were used per extraction, and only those yielding high-quality RNA were used for downstream analysis. RNA concentrations were measured using a NanoDrop spectrophotometer (Thermo Fisher), and RNA integrity was assessed using the Agilent 2100 Bioanalyzer. Mean, range RIN for each group in the study cohort can be found in **Extended data table 1**.

### Sequencing and alignment

All sequencing and alignment steps were performed by Novogene Bioinformatics Technology Co., Ltd. Sample qualities were monitored with RNA integrity numbers using the Bioanalyzer 2100 system (Agilent Technologies, CA, USA). Sequencing libraries were constructed by purifying mRNA from total RNA using poly-T oligo-attached magnetic beads. Purified mRNA was fragmented, and first-strand cDNA was synthesized using random hexamer primers. Second-strand cDNA was synthesized in the presence of dUTP to generate strand-specific libraries, followed by end repair, A-tailing, adapter ligation, size selection, amplification, and purification. Library quality was evaluated with Qubit quantification, real-time PCR, and Bioanalyzer assessment of fragment size distribution. Following library quality control, libraries were pooled according to effective concentration and targeted sequencing depth and sequenced as 150bp paired end reads on an Illumina NovaSeq™ 6000 System. Prior to quality control, an average of 47.5M reads per sample were sequenced. Raw sequencing files (FASTQ) were filtered by Novogene using in-house Perl scripts to remove i) reads with adapter contamination, ii) reads in which undetermined nucleotides constituted >10% of bases, and iii) reads in which low quality bases (Phred quality <5) constituted >50% of the read. After filtering, sequencing data quality was assessed by calculating Q20 (97.32%), Q30 (92.60%), and GC content (50.20%) across samples. Filtered reads were aligned to the human reference genome (hg38) using HISAT2 v2.0.5^59^ with gene annotations from Ensembl (release 94)^60^. An average of 46.4M reads per sample remained, of which 90.63% mapped to hg38 and 87.63% mapped uniquely. Gene-level readcounts were generated using featureCounts v1.5.0-p3^61^ and converted to Fragments Per Kilo base of transcript per Million mapped fragments (FPKM) by Novogene (**Supplementary File 6**).

### Global transcriptomic trend analysis

Prior to global trend analyses, genes with fewer than 10 raw read counts in at least half of the samples were removed to exclude lowly and sparsely expressed transcripts. Separate DESeq2^62^ datasets were then constructed for both the amygdala and frontal cortex from the filtered read counts, with the DESeq2 design formulas including disease and TDP-43 pathological subtyping. We then applied the DESeq2 variance stabilizing transformation (VST) to each tissue-specific dataset (with blind = TRUE) to improve homoscedasticity and normality. Importantly, VST-transformed counts were retained at the individual sample level, and subgroup-level summaries were derived post hoc to produce normalized, averaged expression profiles for visualization purposes. These subgroup-level VST matrices were then used to compute pairwise Euclidean distances between disease and TDP-43 subtype groups, and the resulting distance matrices were visualized for both the amygdala and frontal cortex using the pheatmap package in R^58^.

### Differential expression analysis and visualization

Differential expression (DE) analysis comparing each sample to a tissue-matched cognitively normal control was performed by Novogene Bioinformatics Technology Co., Ltd. using DESeq2 package^62^ in R^58^. The resulting p-values were corrected for multiple testing using the Benjamini-Hochberg method. Differentially expressed genes (DEGs) were defined as those with an adjusted *p*-value < 0.05 and an absolute log_2_ fold change > 0.5 relative to tissue-matched controls. To examine the tissue-wide patterns of differential gene expression across disease and TDP-43 subgroups, we constructed Venn diagrams stratified by tissue (amygdala, frontal cortex) using the ggVennDiagram^63^ package in R. Only genes with the same direction of differential expression in both conditions were counted as overlapping; for example, if a gene measured in the amygdala was significantly upregulated in AD/LATE type α and AD/LATE type β, but was significantly downregulated in AD no TDP, it was counted once in the overlap between AD/LATE type α and AD/LATE type β, and once more for AD no TDP, resulting in a total of two counts for the same gene in the diagram. Volcano plots were constructed using ggplot2^64^ in R to visualize the distribution of log_2_ fold changes and *p* values and to highlight highly differentially expressed genes per subgroup. Within each subgroup-level plot, genes with higher opacity were flagged as “unique” when they were significantly differentially expressed only in the subgroup of interest, and not in any other subtype group, taking direction of regulation into account.

### Biweight midcorrelation analysis of selected differentially expressed genes with pTDP-43 and pTau

Biweight midcorrelation (BiCor) analyses were performed to relate bulk RNA sequencing gene expression to pTDP-43 and pTau protein levels. Gene expression levels were quantified as FPKM (Fragments Per Kilobase of transcript per Million mapped reads) from bulk RNA sequencing of the frontal cortex and amygdala, and pTDP-43 and pTau levels were quantified in absolute values, as described above. For each analysis, we restricted BiCor calculations to genes that were significantly differentially expressed in the disease and TDP-43 subtype group and tissue being investigated. Outliers in both gene expression (FPKM) and protein measurements (pTDP-43 and pTau) falling below the 1st percentile or above the 99th percentile were imputed to NA. All remaining continuous variables were then transformed using a log_10_(x + 1) transformation to reduce skewness and stabilize variance prior to correlation. Biweight midcorrelation coefficients and associated *p*-value were calculated using the bicorAndPvalue function from the WGCNA package^65,66^ in R^58^, treating log_10_(FPKM + 1) for each DEG as the predictor and log_10_(pTDP-43 + 1) or log_10_(pTau + 1) as the outcome. For each gene, we obtained three sets of biweight midcorrelation statistics: (i) correlations with pTDP-43 within the single disease and TDP-43 subtype group of interest, (ii) correlations with pTau within that same subgroup, and (iii) correlations with pTau across all AD groups in the corresponding tissue (AD/LATE type α, AD/LATE type β, and AD no TDP). Genes were considered significantly correlated if they exhibited an absolute biweight midcorrelation coefficient (*r*) ≥ 0.3 and a *p*-value < 0.05. For visualization, we depicted the pTDP-43 correlations from the single-group analyses in volcano-style plots using the ggplot2 package^57^ in R, with the x-axis representing the BiCor coefficient (*r*) and the y-axis representing -log_10_(*p*-value). All genes that were significantly associated with pTDP-43 in the single-group analysis and/or pTau in either the single-group or all-AD analyses were plotted according to their pTDP-43 BiCor statistics, and genes significantly correlated with pTDP-43 but not with pTau (whether evaluated within the specific subtype or across all AD groups) were further highlighted with a black outline. This design addresses the ubiquitous burden of pTau across AD’s various pathological subtypes and therefore allows us to account for genes associated with pTau in both a tissue-wide AD context and a subgroup-specific context, while still restricting pTDP-43 analyses to the TDP-43-stratified groups to maintain sensitivity for subtype-specific associations.

### Gene set pathway enrichment analysis

Gene set enrichment analysis was performed on uniquely differentially expressed genes in each subgroup of interest using ShinyGO (0.82)^67^ with default settings. Enrichment was assessed using the KEGG pathways and GO biological process terms. Pathways were considered significantly enriched if the Benjamini-Hochberg-corrected *p*-value < 0.05. The background transcriptome was the ShinyGO human reference genome.

### Sankey comparison analysis

To identify pathology-specific transcriptional associations in the amygdala across AD/LATE subtypes, genes significantly correlated with quantitative measures of pTDP-43 or pTau pathology were first identified as described in “gene set pathway enrichment analysis”. Pathology-specific gene sets were defined as genes significantly associated with one pathology and not significantly associated with the other pathology (pTDP-43 or pTau). For Sankey diagrams, pathway enrichment analysis was initially performed using the complete set of DEGs prior to application of pathology-specific filters. Enriched GO and KEGG terms were consolidated using REVIGO or KEGG parent pathways to group similar terms. Following consolidation, pathways were mapped to pathology-specific gene sets based on the presence of uniquely correlated genes. Pathways containing genes significantly correlated with both pTDP-43 and pTau were excluded from pathology-specific classification. Relationships between pathology-specific genes and enriched pathways were visualized using Sankey diagrams, illustrating the contribution of pTDP-43-specific and pTau-specific genes to consolidated pathway categories. Need to add Sankey reference for representation.

## Data availability

Raw sequencing data and bam files will be submitted to dbGAP. The datasets generated during the current study are available within the paper of the corresponding author upon reasonable request. Any additional information on the data reported in this paper is available from the lead contact upon request. Source data are provided with this paper.

## Code availability

Previously published software and standard analysis packages were used to perform data analysis as described in Methods.

## Acknowledgements

We thank all the patients and their families for their contribution to this study. This work was supported by the National Institutes of Health (R01NS120992: K.A.J. and M.P., U54NS123743: PF, L.P. and M.P., R01AG037491: K.A.J., P30AG062677 to B.F.B., and U01AG006786 to R.C.P., R35NS097273: L.P.), the Target ALS Foundation (L.P. and M.P.), the BrightFocus Foundation (A2024017S: M.P.), the Kissick Family Foundation (M.P.).

## Author Contributions

X.W., A.C.W., and M.M.R. performed data analysis, prepared data files, and generated figures. J.G.B., Y.S., J.D., M.Y., J.B., C.C., A.N. and M.P. performed RNA and protein extractions, conducted protein immunoassays. R.B.T. and Y.Z. performed histological staining. B.R., E.E.-C., M.D., M.E.M., and D.W.D. processed, evaluated, and provided the postmortem human brain tissue. N.R.G.-R., B.F.B., R.C.P., D.S.K., B.O., G.S.D., and K.A.J. performed neurological evaluations and provided clinical patient data. M.P. conceptualized the study, provided core ideas, and supervised the overall project. L.P., Y.Z., and K.A.J. provided additional project supervision and guidance. X.W., A.C.W., M.M.R., and M.P. wrote and revised the manuscript. All authors reviewed and approved the final manuscript.

## Ethics declarations

Postmortem AD cases from both males and females were obtained from the Mayo Clinic Brain Bank (25-007490), the Mayo Clinic Alzheimer’s Disease Research Center (712-98) and the Mayo Clinic Study of Aging (14-004401). Pathological characterization was performed by an experienced neuropathologist. Written informed consent was given by all participants or authorized family members for the use of brain tissues in research. Study protocols were approved by the institutional review board and ethics committee at Mayo Clinic. Brain tissues (amygdala or frontal cortex) were obtained from 158 individuals with neuropathological diagnosis of AD for total RNA extraction, based on availability.

## Additional information

## Extended data

**Extended Data Table 1. Quantification of pTDP-43 and pTau pathological level across AD and FTLD-TDP cohorts.** The sample median (minimum, maximum) is given for continuous variables, categorical variables are summarized with number and percentage of patients (calculated from total of patients with information in each category).

**Extended Data Fig. 1. Differential gene expression across TDP-43 subtypes in the amygdala and frontal cortex.** Volcano plots showing DEGs across all five study groups in the amygdala (**a–e**) and frontal cortex (**f–j**). DEGs with an absolute log2 fold change > 0.5 and adjusted *p*-value ≤ 0.05 are plotted with high transparency to represent the global magnitude of gene regulation. Genes uniquely dysregulated in each specific group are highlighted with increased opacity. The numbers of total and unique DEGs (upregulated and downregulated) are provided per group in red (upregulated) and blue (downregulated) text in each plot. Uniqueness of genes is defined by both gene identity and direction of regulation. The top 10 genes with the greatest combined score of absolute fold change and statistical significance are labeled.

**Extended Data Fig. 2. Global picture of pathway-level perturbations in all DEGs.** Schematic of pathway-level perturbations in DEGs for each study group. Pink upward and blue downward arrows indicate the directional up- and down-regulation of core biological modules, respectively, across the amygdala (A) and frontal cortex (F). Grey dots indicate that no significant pathway changes were detected in that category.

**Extended Data Fig. 3. Transcriptomic uncoupling of pTau-exclusive signatures in the AD/LATE amygdala.** BiCor volcano plots showing the strength and significance of the correlation between DEG expression and pTau levels. The analyses isolate DEGs that are exclusively correlated with pTau and not pTDP-43. Plots evaluate gene signatures correlated with pTau either exclusively within the specific subtype (single pTau) or globally across all AD samples (all pTau). Panels display these specific correlations in (a) AD/LATE type α with single pTau, (b) AD/LATE type α with all pTau, (c) AD/LATE type β with single pTau, and (d) AD/LATE type β with all pTau. The x-axis represents the biweight midcorrelation coefficient (*r*) for the respective pTau metric, and the y-axis represents -log10(*p*-value). Gene names in blue indicate downregulation; red indicates upregulation. Labeled DEGs are significantly correlated (|*r*| > 0.3, adjusted *p*-value < 0.05).

**Extended Data Fig. 4. Convergence of distinct pTDP-43 and pTau transcriptomic signatures onto shared biological pathways.** Sankey diagrams illustrating pathway-level associations in the amygdala for AD/LATE type α (a-b) and AD/LATE type β (c-d). Condition-specific DEGs containing genes correlating to either pTDP-43 or pTau (a, c) or correlating to pTDP-43, pTau or both (b, d) converge into the same pathways (center nodes). Note pathways were consolidated via REVIGO (detailed in **Supplementary File 5**). Flow thickness represents the number of correlated genes supporting each pathway.

## Supplementary information

**Supplementary File 1. Summary of DEGs and BiCor analysis with pTDP-43 and pTau levels.**

This file contains the complete list of DEGs identified across all disease subtypes (AD/LATE type α, AD/LATE type β, AD no TDP, FTLD-TDP type A, and FTLD-TDP type B) in both the amygdala and frontal cortex. Data includes log_2_ fold changes, *p*-value, and adjusted *p*-value. Additionally, it details the BiCor results linking individual gene expression levels to quantitative biochemical measures of pTDP-43 and pTau pathology, both globally across all AD samples and specifically within individual subtypes.

**Supplementary File 2. KEGG and GO pathway enrichment analysis using all DEGs by condition and tissue.**

This file details the KEGG and GO pathway enrichment analyses performed on the full sets of upregulated and downregulated DEGs for each disease subtype and brain region (amygdala and frontal cortex). Analyses were conducted using a Benjamini-Hochberg-corrected *p*-value cutoff of <0.05. The tables include pathway names, fold enrichment, associated gene counts, and the specific gene lists driving the enrichment for each condition.

**Supplementary File 3. KEGG and GO pathway enrichment analysis using condition- and tissue-specific DEGs.**

This file provides the KEGG and GO pathway enrichment results derived exclusively from unique DEGs, genes whose up- or downregulation is statistically significant and restricted to a single specific disease subtype and/or brain region. By filtering out broadly dysregulated genes, these lists isolate the distinct mechanistic vulnerabilities and specific transcriptomic signatures that define each individual pathology.

**Supplementary File 4. Categorized functional modules of enriched pathways.**

This file aggregates and categorizes the specific pathway enrichment results (from the unique DEGs detailed in **Supplementary File 3**) into major overarching functional biological modules. The tabs categorize the transcriptomic data into specific functional domains including Immune & Inflammation, Metabolism & Bioenergetics, Transcription/Translation & Protein Processing, Cytoskeleton & Cellular Processing, Stress & Stimuli Responses, Vascular & Endothelial, Cell Death, and Neuronal Function, serving as the foundational data for the pathway summaries depicted in **Fig. 3**.

**Supplementary File 5. Pathology-exclusive DEGs and sankey diagram pathway mappings in the AD/LATE amygdala.**

This file focuses on the complex spatial overlap of pathologies in the amygdala of AD/LATE types α and β. It details the BiCor analysis used to identify “exclusive” DEGs, genes whose expression significantly correlates with only pTDP-43 or only pTau. It also includes the data used to generate the Sankey diagrams (**Fig. 4**), detailing how these pathology-exclusive genes map onto parent KEGG and GO pathways (consolidated via REVIGO) to drive specific structural, metabolic, and inflammatory trajectories.

**Supplementary File 6: Quality control metrics for bulk RNA-sequencing.**

This file summarizes the comprehensive quality control metrics for the bulk RNA-sequencing data generated by Novogene. It includes sequencing statistics for all individual patient samples across both the amygdala and frontal cortex. The table details raw and clean read counts, total base counts, overall error rates, Q20 and Q30 quality scores (indicating the percentage of bases with >99% and >99.9% base call accuracy, respectively), and GC content percentages. These metrics validate the robust sequencing depth and high quality of the libraries utilized for all downstream differential expression and transcriptomic analyses.

